# “Tranq-Dope” Overdose and Mortality: Lethality Induced by Fentanyl and Xylazine

**DOI:** 10.1101/2023.09.25.559379

**Authors:** Mark A. Smith, Samantha L. Biancorosso, Jacob D. Camp, Salome H. Hailu, Alexandra N. Johansen, Mackenzie H. Morris, Hannah N. Carlson

**Affiliations:** Department of Psychology and Program in Neuroscience, Davidson College, Davidson, NC, USA

**Keywords:** addiction, opioid use disorder, preclinical model, sex differences, substance use, substance use disorder

## Abstract

The recreational use of fentanyl in combination with xylazine (i.e., “tranq-dope”) represents a rapidly emerging public health threat characterized by significant toxicity and mortality. This study quantified the interactions between these drugs on lethality and examined the effectiveness of potential rescue medications to prevent a lethal overdose. Male and female mice were administered acute doses of fentanyl, xylazine, or their combination via intraperitoneal injection, and lethality was determined 30, 60, 90, 120, and 1440 min (24 hr) after administration. Both fentanyl and xylazine produced dose-dependent increases in lethality when administered alone. A nonlethal dose of fentanyl (56 mg/kg) produced an approximately 5-fold decrease in the estimated LD_50_ for xylazine (i.e., the dose estimated to produce lethality in 50% of the population). Notably, a nonlethal dose of xylazine (100 mg/kg) produced an approximately 100-fold decrease in the estimated LD_50_ for fentanyl. The opioid receptor antagonist, naloxone (3 mg/kg), but not the alpha-2 adrenergic receptor antagonist, yohimbine (3 mg/kg), significantly decreased the lethality of a fentanyl-xylazine combination. Lethality was rapid, with death occurring within 10 min after a high dose combination and generally within 30 min at lower dose combinations. Males were more sensitive to the lethal effects of fentanyl-xylazine combinations under some conditions, suggesting biologically relevant sex differences in sensitivity to fentanyl-xylazine lethality. These data provide the first quantification of the lethal effects of “tranq-dope” and suggest that rapid administration of naloxone may be effective at preventing death following overdose.

## 1. Introduction

Opioid misuse and addiction represent an ongoing public health crisis. In the United States, the annual number of deaths attributed to opioids now exceeds the annual number of deaths from firearms and motor vehicle accidents (Center for Disease Control, 2023b). Because nonlethal overdoses are much more common than lethal overdoses, annual mortality rates significantly underestimate the proportion of the population who experience clinically significant but nonlethal consequences of an acute overdose (Center for Disease Control, 2023a). Complicating matters further, ∼80% of opioid overdoses in the United States are characterized by polysubstance use involving a nonopioid drug (Jones et al., 2018). Many of these drugs increase the lethal effects of opioids, making lethal overdoses more common and less sensitive to reversal by traditional rescue medications.

The fully synthetic mu opioid receptor agonist, fentanyl, has driven the most recent wave of the opioid overdose epidemic, partly because of its increased availability and greater potency relative to heroin (Gladden, 2016; O’Donnell, 2017). Between 2019 and 2022, the monthly percentage of fentanyl-related overdose deaths increased across 21 jurisdictions in the United States by over 200% (Kariisa et al., 2023). Concomitant with this increase in fentanyl-related mortality, seizures of xylazine-adulterated fentanyl (“tranq-dope”) increased by a similar amount across many of these jurisdictions (United States Drug Enforcement Agency, 2023). Xylazine is an alpha-2 adrenergic receptor agonist used as a general anesthetic in veterinary medicine but not approved for use in humans. Both the Center for Disease Control and the US Drug Enforcement Administration have identified fentanyl-xylazine combinations as a significant public health threat (Center for Disease Control, 2023c; United States Drug Enforcement Agency, 2023), and reports of the human impact of this deadly drug combination has been described in both scientific (Gipson and Strickland, 2023) and lay (Hoffman, 2023) publications. The rapid increases in lethal drug overdoses attributed to this drug combination recently prompted the White House Office of National Drug Control Policy to declare xylazine combined with fentanyl an emerging threat to the United States (Office of National Drug Control Policy, 2023). Although evidence suggesting xylazine increases fentanyl-related morbidity and mortality is rapidly growing, no studies have quantified the interactions between these drugs on lethality.

The purpose of this study was to quantify the lethal effects of fentanyl and xylazine, alone and in combination with one another, in male and female mice. To this end, acute doses of fentanyl and xylazine were administered alone or in various dose combinations, and lethality was determined at several time points over the course of 24 hours. Lethality was quantified by calculating estimated LD_20_, LD_50_, andLD_80_ values (i.e., the dose estimated to produce lethality in 20%, 50%, and 80% of the population, respectively) for each drug alone and in combination with one another. Additional tests were conducted to determine whether a mu opioid receptor antagonist (naloxone) and an alpha-2 receptor antagonist (yohimbine), both of which are available for use in humans, could prevent fentanyl-xylazine lethality when administered immediately after a potentially lethal dose combination. Finally, tests were conducted to measure the time course of lethality following a high dose combination of fentanyl and xylazine, and biologically relevant sex differences in sensitivity to fentanyl-xylazine lethality were determined.

## 2. Materials and Methods

### 2.1. Subjects

Male and female C57BL/6 mice were obtained at 42 days of age from Charles River Laboratories (Raleigh, NC, USA) and housed in same-sex groups of five mice. Food, water, and enrichment items were continuously present in the home cage. Mice habituated to the vivarium for a minimum of five days before testing. All procedures were approved by the Davidson College Institutional Animal Care and Use Committee.

### 2.2. Procedure

A total of 347 mice (male = 172; female = 175) were used in this study. All tests were conducted in 6 to 18 mice, with equal numbers of males and females represented at each dose or dose combination, except in two instances in which the number of males and females differed by no more than two mice.

Initial doses of xylazine and fentanyl were selected for testing based on a literature search of their potency to produce maximal effects on measures of analgesia and anesthesia in rodents, and then moving in quarter log units upward or downward until at least four doses were tested, with at least one dose producing greater than 80% lethality and at least one dose producing less than 20% lethality. For drug combination tests, doses of fentanyl (56 mg/kg) and xylazine (100 mg/kg) were selected that represented the lowest dose of each drug that did not produce lethality (0% lethality; 100% survival) when tested alone.

On the day of testing, mice were transferred to individual cages with food, water, and bedding. Approximately one hour after transfer, mice were randomized and injected intraperitoneally with fentanyl, xylazine, or a combination of fentanyl and xylazine. Mice were checked for lethality 30, 60, 90, 120, and 1440 min (24 hr) after administration by unblinded observers. Lethality was operationally defined as the absence of (1) movement, (2) respiration, and (3) heartbeat. Any mouse remaining alive after 24 hr was euthanized according to procedures approved by the American Veterinary Medical Association.

Rescue tests measuring prevention of lethality produced by 32 mg/kg fentanyl + 56 mg/kg xylazine were conducted with 3 mg/kg naloxone, 3 mg/kg yohimbine, and 3 mg/kg naloxone + 3 mg/kg yohimbine. In these tests, a single intraperitoneal injection of naloxone, yohimbine, or a combination of naloxone and yohimbine was given immediately following the fentanyl-xylazine dose combination. Lethality was then measured as described above.

To characterize the time course of lethality, we determined, *a priori*, that lethality checks would be scheduled 30, 60, 90, 120, and 1440 min (24 hr) after administration; however, in the majority of instances, lethality occurred within the first 30 min. Consequently, we conducted an additional time-course test in which lethality was measured at 2-min intervals, beginning immediately after administration. In this test, we administered a dose combination of 56 mg/kg fentanyl + 100 mg/kg xylazine, doses that were not lethal when administered individually, but produced 100% lethality when administered in combination during pilot testing. In this time-course test, we measured lethality as described above every 2 min until lethality was observed.

### 2.3. Drugs

Fentanyl HCl was generously supplied by the National Institute on Drug Abuse (Research Triangle Institute, Research Triangle Park, NC, USA). Naloxone HCl, xylazine HCl, and yohimbine HCl were purchased from Sigma Chemical Co. (St. Louis, MO, USA). All drugs were dissolved in sterile saline and administered via intraperitoneal injection in a volume of 10 ml/kg.

### 2.4. Data Analysis

The primary dependent measure of this study was the percentage of mice dead 24 hr after drug administration (% lethality). These data were used to determine the estimated lethal dose (LD) of fentanyl, xylazine, and fentanyl-xylazine combinations. Estimated LD_1_ to LD_99_ values and their 95% confidence limits (95% CL) were determined via probit analysis (Finney, 1952) using an open-source calculator (Mekapogu, 2021). Differences between estimated LD_50_ values served as the primary outcome variable when comparing lethality across conditions, with differences in estimated LD_20_ and LD_80_ values serving as secondary outcome variables. Estimated LD values were considered statistically significant when 95% confidence limits did not overlap.

Data from rescue tests were analyzed via a χ^2^ test using % lethality obtained in males and females after 24 hours. This analysis was not powered to detect differences across conditions within each sex.

Latency to produce lethality was determined visually using heat maps depicting the % lethality at each observed time point after drug administration.

All data are available with no restrictions at Mendeley Data (Smith, 2023).

## 3. Results

### 3.1. Fentanyl-Xylazine Lethality

Xylazine produced dose-dependent increases in lethality characterized by a steep dose-effect curve in which 0% to 100% lethality was observed within 0.5 log unit (Figure 1). Lethality was rapid, with death occurring within 30 min in all instances in which lethality was observed (Supplemental Figure 1). A dose of fentanyl representing the highest dose that did not produce lethality when administered alone (56 mg/kg) shifted the xylazine dose-effect curve to the left. The estimated LD_50_ value (95% CL) of xylazine was 157.2 mg/kg (138.2 - 178.7 mg/kg) alone versus 32.0 mg/kg (23.2 - 44.1 mg/kg) in the presence of fentanyl, representing a significant 4.9-fold increase in potency (Table 1). This dose of fentanyl also flattened the xylazine dose-effect curve as revealed by greater decreases in its estimated LD_20_ value relative to its estimated LD_80_ value (8.5-vs. 2.9-fold increases in potency, respectively). Lethality was not as rapid at lower unit doses of xylazine in the presence of fentanyl, with death occurring after 60 min in some mice (Supplemental Figure 1).

**Figure 1.**
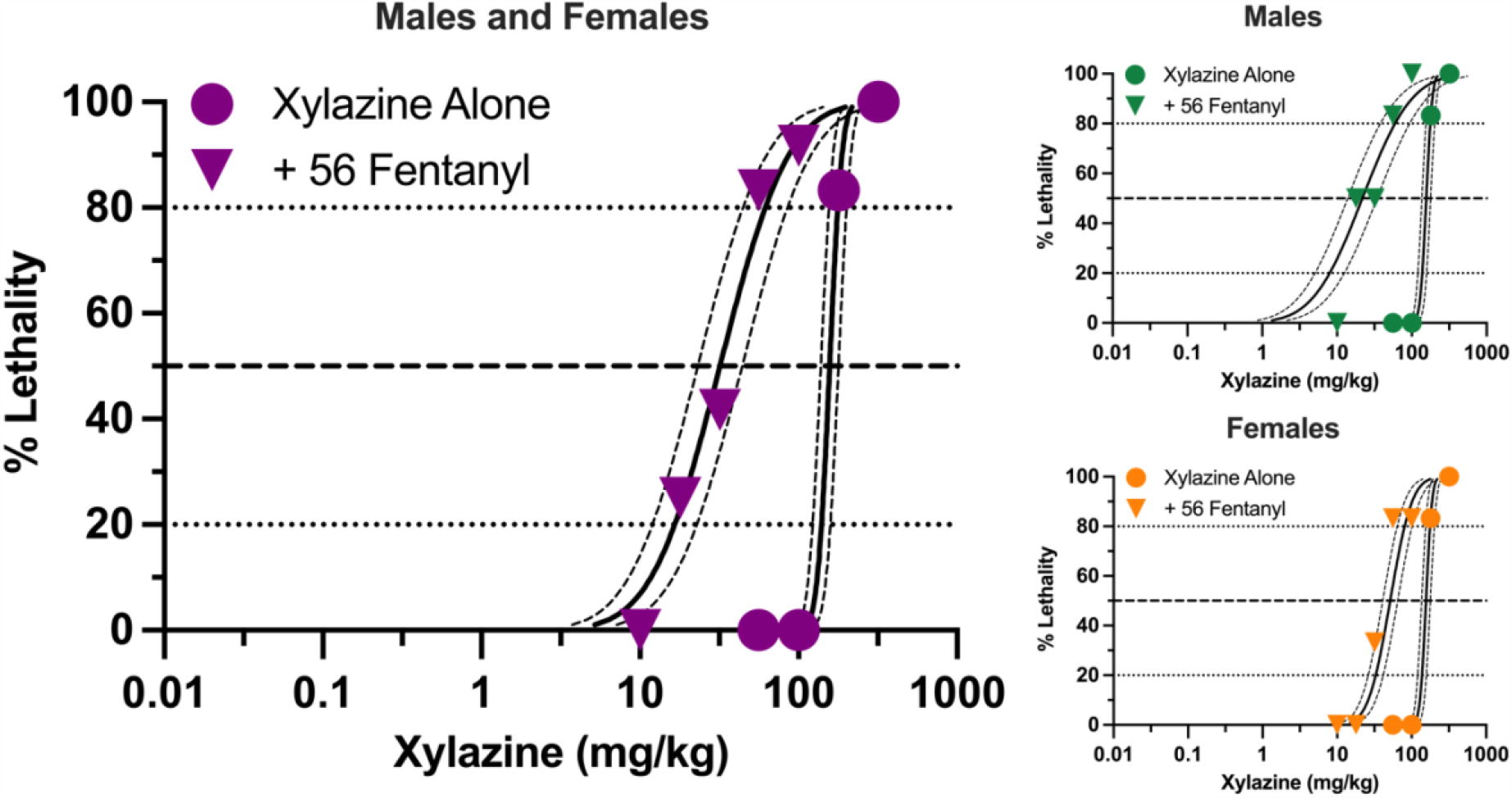
Lethal effects of xylazine alone and in combination with 56 mg/kg fentanyl in male and female mice. Left: Lethality produced by various doses of xylazine alone (circles) and in combination with 56 mg/kg fentanyl (triangles). Left axis depicts percentage (%) of mice dead after 24 hours. Bottom axis depicts doses of xylazine in mg/kg. All points represent data obtained from 6-18 mice. Spline curves represent curves of best fit (95% CL) for points representing the estimated LD_1_ – LD_99_ for xylazine alone and xylazine in combination with 56 mg/kg fentanyl. Upper Right: Lethal effects of xylazine alone and in combination with 56 mg/kg fentanyl in male mice. All points represent data from 3 to 9 mice. Lower Right: Lethal effects of xylazine alone and in combination with 56 mg/kg fentanyl in female mice. All data points represent data from 3 to 9 mice.

Males and females did not differ in sensitivity to xylazine-induced lethality when it was administered alone (Figure 1), and estimated LD_20_, LD_50_, and LD_80_ values were identical between the two sexes (Table 1). Males were more sensitive than females to xylazine in the presence of 56 mg/kg fentanyl as revealed by significantly lower estimated LD_20_ and LD_50_, values; however, these differences were relatively small, and estimated LD values only differed 3-fold between males and females at their most extreme.

Similar to xylazine, fentanyl produced dose-dependent increases in lethality characterized by a steep dose-effect curve in which 0% to 100% lethality was observed within 0.75 log unit (Figure 2). Lethality was not as rapid as that observed with xylazine alone, with some rats dying more than 60 min after administration (Supplement Figure 2). A dose of xylazine representing the highest dose that did not produce lethality (100 mg/kg) shifted the fentanyl dose-effect curve markedly to the left. The estimated LD_50_ value (95% CL) of fentanyl was 131.3 mg/kg (107.8 - 159.9 mg/kg) alone versus 1.27 mg/kg (0.74 - 2.16 mg/kg) in the presence of xylazine, representing a significant, 103.3-fold increase in potency (Table 1). This dose of xylazine also flattened the fentanyl dose-effect curve as revealed by greater decreases in its estimated LD_20_ value relative to its LD_80_ value (381.9-vs. 28.0-fold increases in potency, respectively). Lethality was rapid at all dose combinations, occurring in less than 30 min on all but one occasion in which death occurred (Supplemental Figure 2). Males were slightly more sensitive than females to fentanyl-induced lethality when administered alone and in combination with xylazine (Figure 2); however, these differences were not significant, and confidence limits for all estimated LD values for males overlapped those for females (Table 1).

**Figure 2.**
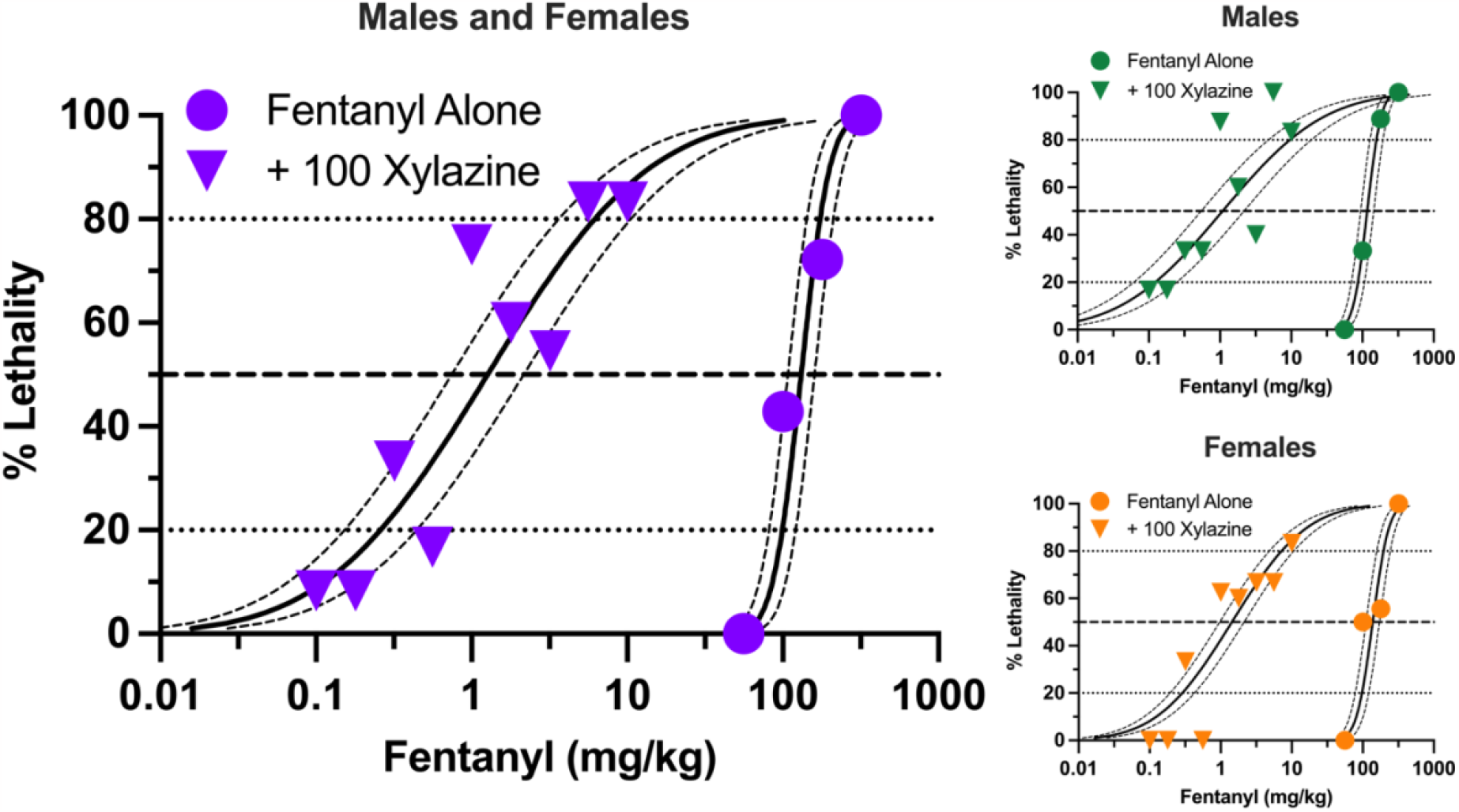
Lethal effects of fentanyl alone and in combination with 100 mg/kg xylazine in male and female mice. Left: Lethality produced by various doses of fentanyl alone (circles) and in combination with 100 mg/kg xylazine (triangles). All points represent data obtained from 6-16 mice. All other details are as described in Figure 1. Upper Right: Lethal effects of fentanyl alone and in combination with 100 mg/kg xylazine in male mice. All points represent data obtained from 3-8 mice. Lower Right: Lethal effects of fentanyl alone and in combination with 100 mg/kg xylazine in female mice. All points represent data obtained from 3-8 mice.

### 3.2. Effects of Potential Rescue Medications

We next determined the effectiveness of the opioid receptor antagonist, naloxone, and the alpha-2 adrenergic receptor antagonist, yohimbine, to prevent fentanyl-xylazine lethality. To this end, mice were administered a combination of 32 mg/kg fentanyl and 56 mg/kg xylazine, doses that were 0.5 log unit lower than those required to produce lethality when administered alone. Mice were then immediately administered a second injection of either 3 mg/kg naloxone, 3 mg/kg yohimbine, or a combination of 3 mg/kg naloxone + 3 mg/kg yohimbine. The dose combination of fentanyl + xylazine produced 60% lethality in the absence of any rescue medication (Figure 3), with most mice dying in the first 30 min (Supplemental Figure 3). Naloxone significantly decreased the lethality of this dose combination to approximately 20%, which was significantly different from both vehicle (χ^2^ = 5.60, *p* = .018, *d* = 0.86) and yohimbine (χ^2^ = 9.47, *p* = .002, *d* = 1.24). Lethality was not altered by yohimbine. Naloxone in combination with yohimbine marginally reduced lethality of fentanyl + xylazine, but this effect was not statistically significant. No sex differences were observed for any drug condition. The latency to lethality varied across mice, occurring greater than 60 min after administration in a small percentage of mice (Supplemental Figure 3). Rescue tests were not powered to detect differences across drug conditions within each sex; however, patterns of lethality were very similar between males and females (Figure 3).

**Figure 3.**
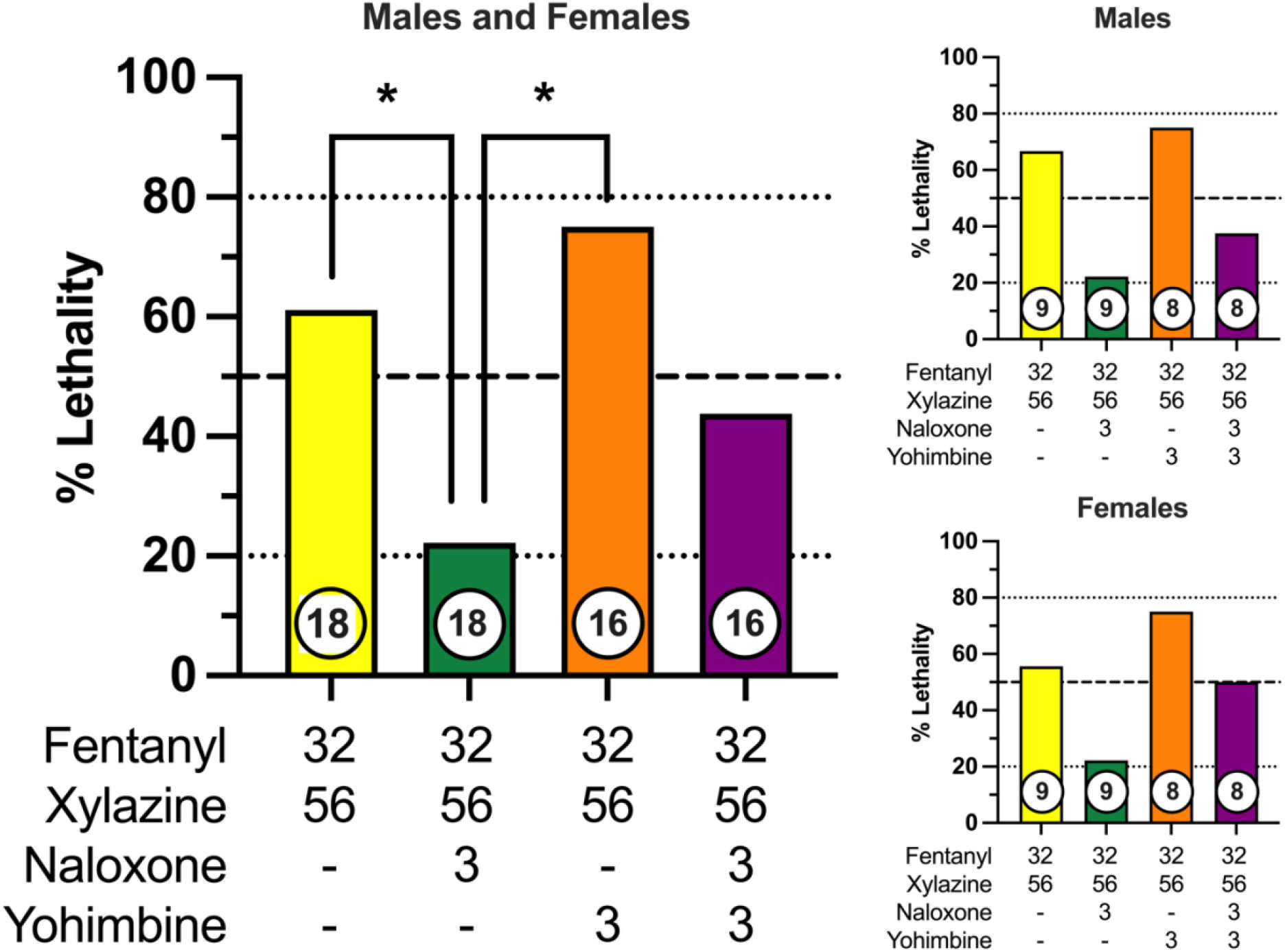
Prevention of fentanyl/xylazine lethality by naloxone and/or yohimbine in male and female mice. Left: Lethality produced by 32 mg/kg fentanyl + 56 mg/kg xylazine in combination with vehicle, 3 mg/kg naloxone, 3 mg/kg yohimbine, and 3 mg/kg naloxone + 3 mg/kg yohimbine. Left axis depicts percentage (%) of mice dead after 24 hours. Bottom axis depicts doses of fentanyl, xylazine, naloxone, and yohimbine. Numbers in circles indicate number of mice tested. Asterisks (*) indicate significant differences between groups (*p* < .05) . Upper Right: Prevention of fentanyl/xylazine lethality by naloxone and/or yohimbine in male mice. Lower Right: Prevention of fentanyl/xylazine lethality by naloxone and/or yohimbine in female mice.

### 3.3. Lethality Time Course

The primary outcome measure of this study was % lethality 24 hr after fentanyl-xylazine administration. To characterize the time course of lethality, we determined, *a priori*, that lethality checks would be scheduled 30, 60, 90, 120, and 1440 min after administration; however, in the majority of instances, lethality occurred within the first 30 min. Specifically, in the 116 instances in which a mouse died, lethality was observed within the first 30 min on 100 occasions (86.2%). Consequently, we conducted an additional test in which lethality was measured at 2-min intervals, beginning immediately after administration. To this end, we administered a dose combination of 56 mg/kg fentanyl + 100 mg/kg xylazine, doses that were not lethal when administered alone, to 8 mice (4 male, 4 female). This dose combination produced 100% lethality within 10 min of administration (Figure 4), and males died significantly faster than females [*t*(6) = 4.02, *p* = .007, *d* = 2.85]. Notably, all males died within 2-6 min, whereas all females died between 8-10 min, suggesting males may be more vulnerable to the effects of a high dose combination.

**Figure 4.**
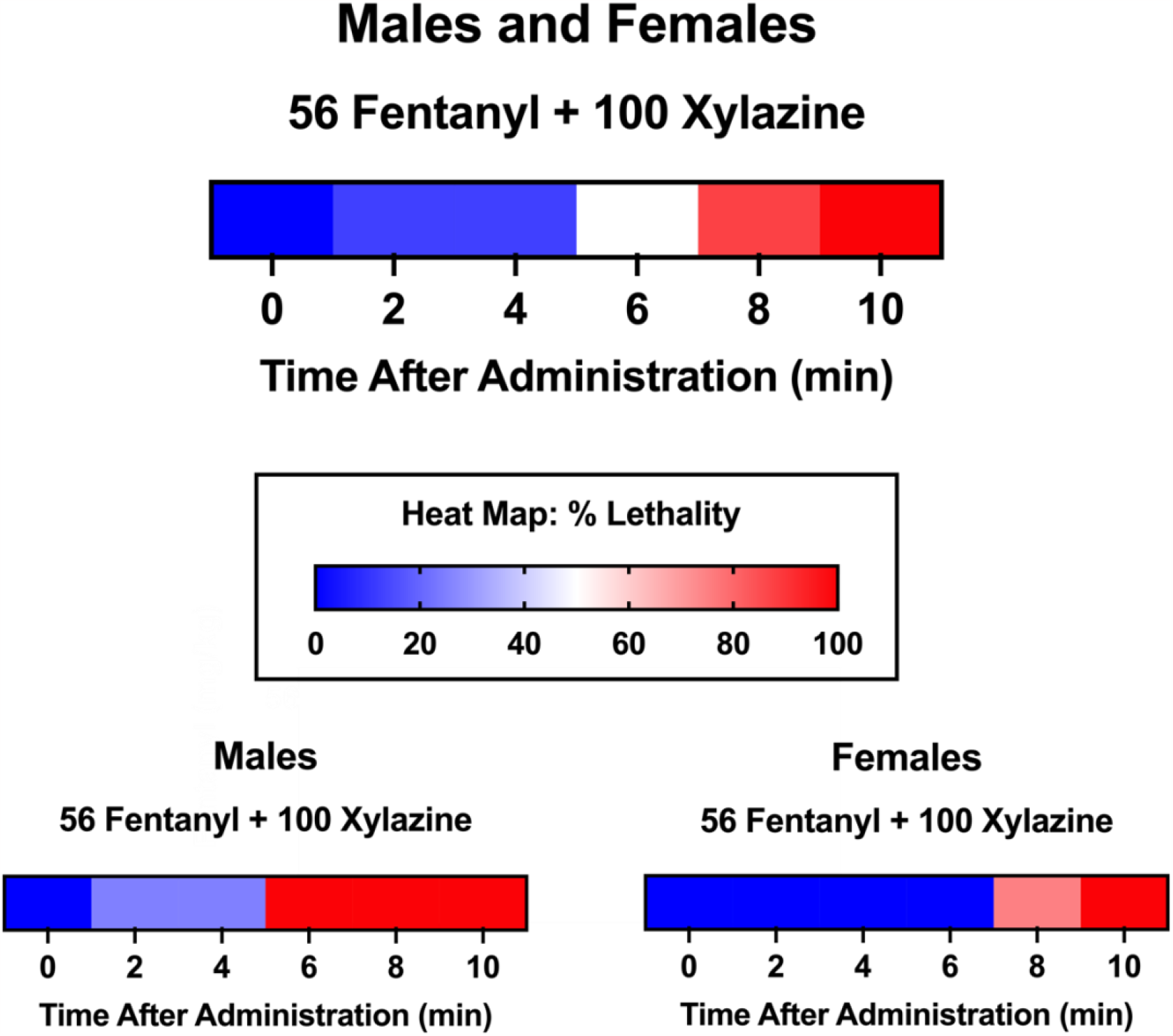
Heat maps depicting time course of lethality produced by 56 mg/kg fentanyl + 100 xylazine. Lethality was determined 2, 4, 6, 8, and 10 min after administration. Color depicts % lethality from 0 (dark blue) to 100 (dark red). Data are depicted for male and female mice (top; n = 8), male mice only (lower left; n = 4), and female mice only (lower right; n = 4).

## 4. Discussion

The principal findings of this study are that (1) fentanyl and xylazine each produce lethality characterized by steep dose-effect curves, (2) fentanyl and xylazine significantly increase the lethal effects of one another, often to very large degrees, (3) lethality produced by this drug combination is generally more rapid than high doses of fentanyl administered alone, (4) naloxone significantly decreases fentanyl-xylazine lethality when administered immediately after exposure, (5) yohimbine does not alter fentanyl-xylazine lethality and does not increase (and may decrease) the effectiveness of naloxone, and (6) males are more sensitive than females to fentanyl-xylazine lethality under some conditions.

The ability of fentanyl and xylazine to increase the lethality of the other was substantial. A nonlethal dose of fentanyl shifted the midpoint of the xylazine lethality curve 5-fold to the left, meaning that the median lethal dose of xylazine was 5-fold smaller in the presence of fentanyl. Notably, this shift was greater at lower doses of xylazine due to the nonparallel shift in the dose-effect curve. A markedly greater shift was observed when a nonlethal dose of xylazine was combined with fentanyl. This dose of xylazine shifted the midpoint of the fentanyl lethality curve ∼100 fold to the left, representing an increase in potency of two orders of magnitude. Again, this increase in potency was greatest for the most sensitive quintile, with the LD_20_ shifting ∼380 fold to the left. The magnitude of this potency increase represents a significant public health threat when considered at a population level and is largely unprecedented.

The opioid receptor antagonist, naloxone, is the primary rescue medication for suspected opioid overdose; however, naloxone is less effective against fentanyl than heroin (Pergolizzi et al., 2021), and similar reports have noted its reduced effectiveness for fentanyl-xylazine combinations (Pergolizzi et al., 2023). The dose of naloxone used in this study (3 mg/kg) is fully effective at reversing fentanyl-induced respiratory depression in fentanyl-treated mice (Hill et al., 2020). In the present study, this dose of naloxone was significantly but only partially effective at preventing fentanyl-xylazine lethality, despite using doses of fentanyl and xylazine that were 0.5 log unit lower (i.e., ∼3.2 fold lower) than those required to produce lethality when administered alone. The alpha-2 adrenergic receptor antagonist, yohimbine, is used in veterinary medicine to reverse xylazine anesthesia, and it was selected for this study as a potential rescue medication because it is available for use in humans, due to its legacy as a treatment for erectile dysfunction. Unfortunately, yohimbine showed no evidence of effectiveness at preventing fentanyl-xylazine lethality and may have attenuated the effectiveness of naloxone when administered as part of a combination treatment. We do not know why yohimbine was ineffective or may have reduced the effectiveness of naloxone. Yohimbine is known to produce hypertension, tachycardia, and cardiac arrhythmias (Cimolai and Cimolai, 2011), and this may have contributed to its failure to prevent lethality in animals that were pharmacologically compromised by fentanyl; however, this is speculative and further studies are warranted.

Lethality was rapid in almost all cases. We initially determined lethality 30, 60, 90, 120, and 1440 min after administration to characterize the time course. In the majority of instances in which lethality was observed, it occurred within 30 min. An additional time-time course test in which lethality was determined at 2-min intervals using high doses of fentanyl and xylazine produced lethality in all mice within 10 min. This time course is concerning given the combination was administered via intraperitoneal injection, whereas human populations typically self-administer this combination intravenously. Given that pharmacokinetic parameters related to onset of action and time to peak plasma/tissue concentrations are an order of magnitude faster for intravenous versus intraperitoneal administration (Zhao et al., 2012), these data imply that rescue medications may be needed within 60 s of an overdose to be effective.

Age-adjusted rates of drug overdose deaths for males are approximately twice those for females, and these differences have been apparent throughout the opioid epidemic (Center for Disease Control, 2022). Males and females were equally represented in this study to characterize biological sex differences in sensitivity to fentanyl-xylazine lethality and pharmacological rescue. Findings were generally similar between sexes, but males were more sensitive than females on some measures. Specifically, males were more sensitive to fentanyl-induced leftward shifts in the xylazine dose-effect curve, but this shift was only ∼3-fold greater than that observed in females at its most extreme. Lethality was more rapid in males than females following a high dose combination, with all males and no females dying within the first 6 min of administration. Although naloxone was equally effective in males and females at preventing lethality, these data suggest that the time frame for reversing an overdose may be significantly shorter for males than females.

Strengths of the study include the determination of full dose-response curves for fentanyl and xylazine lethality, the inclusion of both males and females, the inclusion of rescue tests by readily available pharmacological antagonists, and the inclusion of time course determinations in all tests of lethality. Important limitations of the study include the single dose design of the rescue tests and the single dose of fentanyl (56 mg/kg) and xylazine (100 mg/kg) used in the drug-combination experiments to determine changes in LD values. Testing additional doses of rescue medications is particularly important because fentanyl is 100 times more potent than heroin, and naloxone doses currently in use may be below threshold to show efficacy (especially in the new context of xylazine). This is translationally important because empirical evidence is needed to support medications development of more potent naloxone analogues, or at the least, higher doses of naloxone. Another limitation is the absence of intravenous testing, which is the typical route of administration for individuals who inject drugs nonmedically. We selected the intraperitoneal route because of practical concerns related to determining time course measurements across large numbers of animals, as well as increasing the likelihood that rescue medications would be capable of preventing overdose if they were effective. Finally, all tests were conducted in naïve animals, which limits the translational appeal of these data for human populations with an extensive pharmacological history and hence a potential for tolerance.

Data from this study address the call from the White House to mobilize a National Response Plan to produce “a 15% reduction (compared to 2022 as the baseline year) of xylazine positive drug poisoning deaths in at least three of four U.S. census regions by 2025” (The White House, 2023). Findings derived from this study suggest that widespread screening for xylazine in seizures by law enforcement and in clinical samples obtained by emergency departments are needed to better quantify the relative frequency in which street-level fentanyl is adulterated with xylazine, as well as the concentrations of xylazine relative to fentanyl in these adulterated products. Finally, these data also support the need for the widespread availability of high-dose formulations of naloxone.

## Author Contributions

MAS designed the study and wrote the first draft of the manuscript. SLB, JDC, SHH, ANJ, MHM, and HNC collected all data, compiled all data, and contributed to the final draft of the manuscript. All authors approved the final manuscript.

## Acknowledgements

The authors thank Drs. William Fantegrossi and Drake Morgan for comments on a previous version of this manuscript.

