## Supplementary figures and images for "“Tranq-Dope” Overdose and Mortality: Lethality Induced by Fentanyl and Xylazine"

### Supplemental Figure 1

**Supplemental Figure 1**

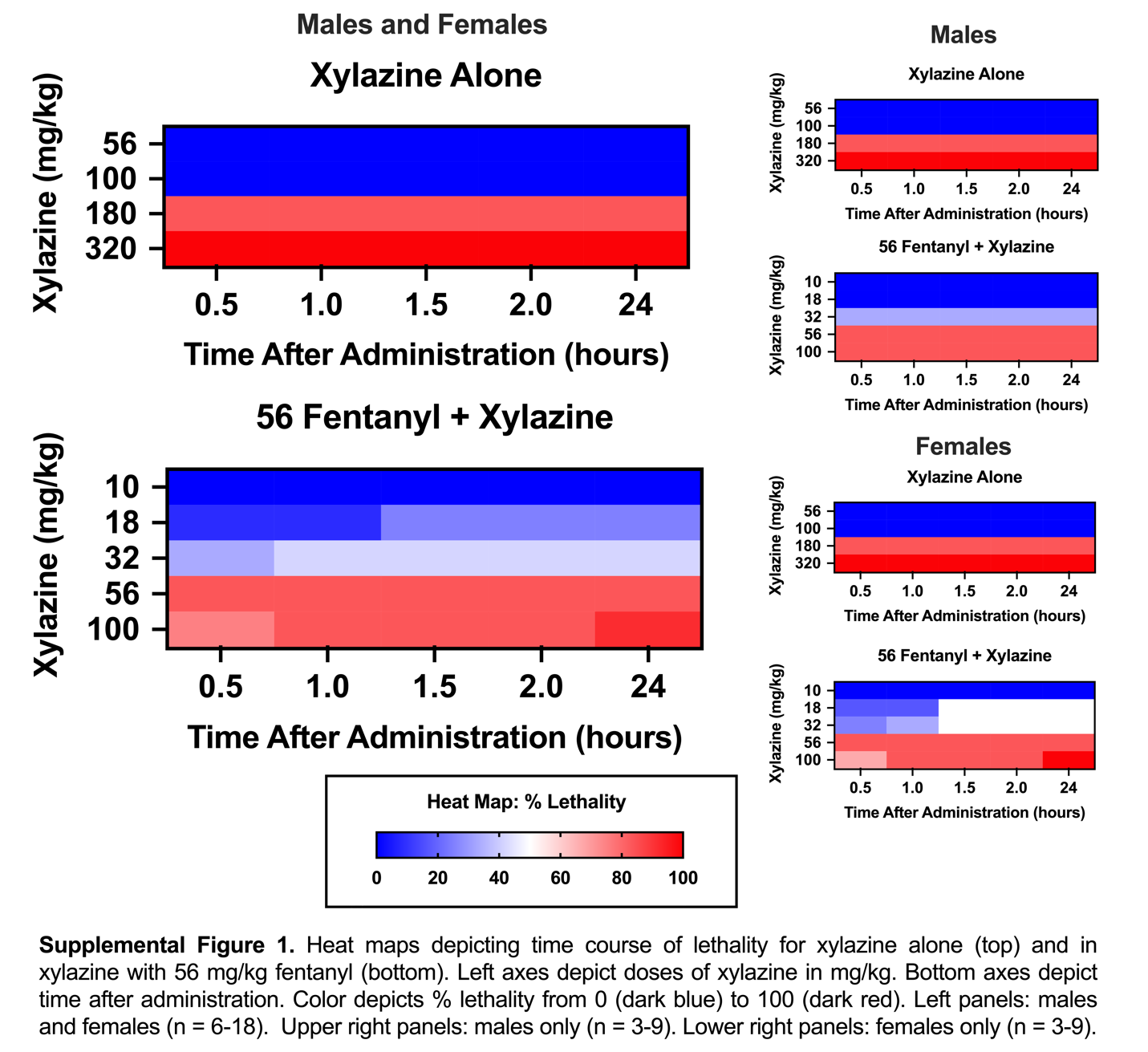

### Supplemental Figure 2

**Supplemental Figure 2**

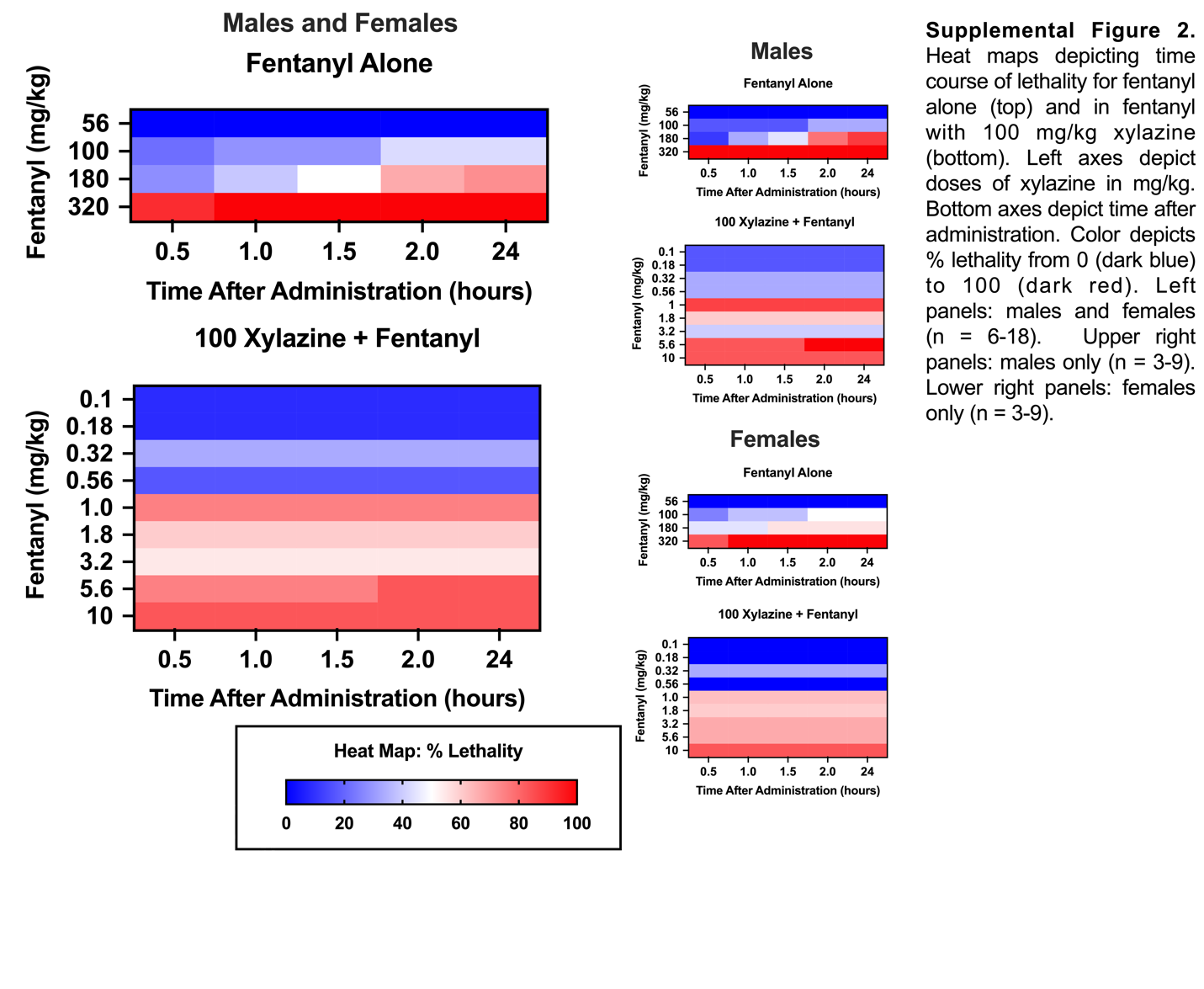

### Supplemental Figure 3

**Supplemental Figure 3**

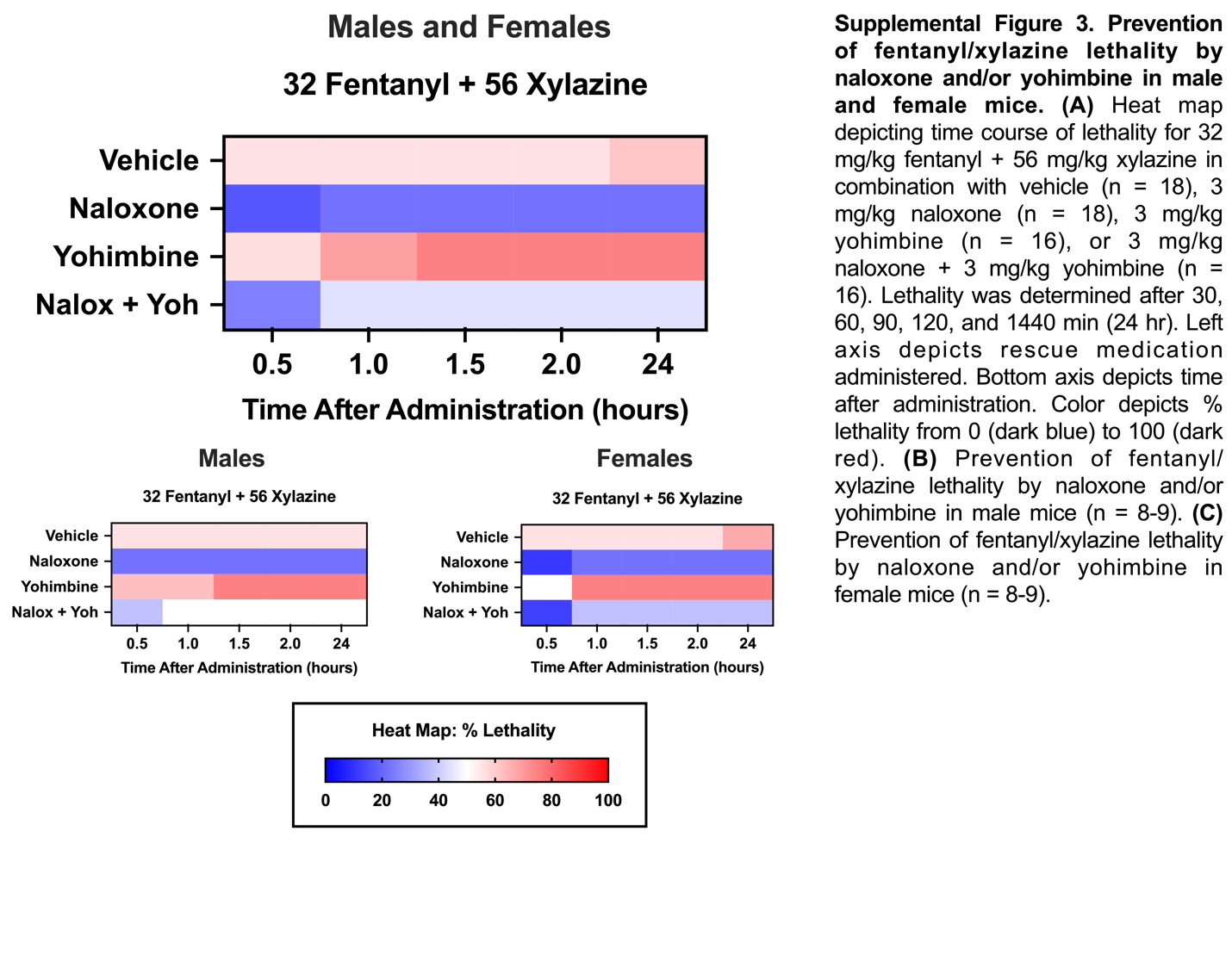
